# Alpha Oscillations Prior to Encoding Preferentially Modulate Memory Consolidation during Wake Relative to Sleep

**DOI:** 10.1101/202176

**Authors:** Zachariah R. Cross, Amanda Santamaria, Andrew W. Corcoran, Phillip M. Alday, Scott Coussens, Mark J. Kohler

## Abstract

Sleep promotes memory consolidation through unique neuromodulatory activity. However, little is known about the impact of attention during pre-sleep memory encoding on later memory performance. The current study aimed to address the question of whether attentional state prior to encoding, as indexed by alpha oscillatory activity, modulates the consolidation of images across periods of sleep and wake. 22 participants aged 18 – 41 years (mean age = 27.3) viewed 120 emotionally valenced images (positive, negative, neutral) before a 2hr afternoon sleep opportunity and an equivalent period of wake. Following the sleep and wake conditions, participants were required to distinguish between 120 previously seen (target) images and 120 new (distractor) images. Relative alpha power – adjusted according to participants’ individual alpha frequency – was computed to index attentional state prior to the learning phase. Generalised linear mixed-effects modelling revealed memory performance was modulated by attention, such that greater pre-encoding alpha power preferentially promoted memory consolidation during wake compared to sleep. There was no difference in memory performance between positive, negative and neutral stimuli. Modulations in alpha oscillatory activity may help to coordinate the flow of information between task-relevant cortical regions and a thalamo-cortical loop that preferentially subserves the formation of memory during times of wake relative to sleep.

## 1. Introduction

The central nervous system appears to prioritise emotional over neutral information. This prioritisation of emotional information modulates memory across encoding, consolidation and retrieval (LaBar & Cabeza, 2006). Encoding is the acquisition of new information in newly constructed representations, while consolidation transforms these representations into stable forms, integrating them into existing knowledge networks, which are then accessible during retrieval (Diekelmann, Wilhelm & Born, 2009). Behavioural (Kensinger, Garoff-Eaton & Schacter, 2007) and neuroimaging (Uribe, Garcia & Tomaz, 2011) studies of emotional memory also reveal that attention (Brenner, Rumak, Burns & Kieffaber, 2014; Maddox, Naveh-Benjamin, Old & Kilb, 2012) and sleep (Groch, Wilhelm, Diekelmann & Born, 2013) are important modulators at encoding and consolidation. At encoding, attention is preferentially allocated to emotional over neutral information, facilitating enhanced consolidation of emotional information (Vuilleumier, 2005). Similarly, it has been suggested that sleep provides a suitable environment for unique neuromodulatory processes to facilitate the distribution of hippocampally-dependent information into neocortical long-term memory (LTM) networks (Ellenbogen, Hulbert, Stickgold, Dinges & Thompson-Schill, 2006). However, little is known regarding the interactive effect of attention and sleep on the emotional modulation of LTM.

### 1.1. Impact of attention on emotional memory

Emotional content influences attention during encoding (Barnacle, Montaldi, Talmi & Sommer, 2016). Regions of the brain activated by emotional content, such as the amygdala, promote hippocampal-dependent memory consolidation (LaBar & Cabeza, 2006; Murty, Ritchey, Adcock & LaBar, 2010; Smith, Henson, Dolan & Rugg, 2004). Similarly, behavioural studies have demonstrated greater attention is paid to emotional compared to neutral content within divided attention tasks, resulting in superior memory for emotional over neutral stimuli at delayed recall intervals (Maddox et al., 2012; Sakaki, Niki & Mather, 2012).

Given the high temporal resolution of electroencephalography (EEG), event related potentials (ERP) are useful in assessing the time course of neural activation patterns of attention to emotional stimuli (for review: Schupp, Flaisch, Stockburger & Junghofer, 2006). However, ERPs are limited in their representation of the neural mechanisms that underpin fluctuations in attention. The excitation of cortical structures may be mediated by rhythmic activity over extended periods of time, reflecting the organisation and interconnectivity of attentional networks (Payne & Sekular, 2014; Pfurtscheller, Stancak & Neuper, 1996; Yordanova, Kolev & Polich, 2001). High amplitude, low frequency oscillations occurring in the alpha-band (~7 – 13 Hz) are believed to index coherent activity in attentional networks (Güntekin & Başar, 2014; Klimesch, 2012; Uusberg, Uibo, Kreegipuu & Allik, 2013).

Alpha desynchronisation (i.e. decrease in amplitude) is viewed as reflecting the activation of cortical areas with increased neuronal excitability, whereas alpha synchronisation (i.e. increase in amplitude) reflects the inhibition of brain regions (Jensen & Mazaheri, 2010; Klimesch, 2012). This inhibitory account of alpha is supported by a series of recent studies (for review: Sadaghiani & Kleinschmidt, 2016), and highlights the need to move beyond the view that alpha simply reflects passive neural idling (cf. Pfurtscheller et al., 1996). For example, Sadaghiani and Kleinschmidt (2016) posit that the regulation of alpha power occurs via a cingulo-opercular network, while dorsal fronto-parietal regions downscale alpha power via a top-down influence on local, task-relevant regions. In support of this view, Walz and colleagues (2015) reported an association between greater alpha desynchronisation and the activation of brain regions involved in attentional processing (i.e., bilateral thalamus and posterior parietal cortex) with enhanced behavioural performance on an auditory oddball task. These findings support proposals (e.g. Hanslymayr et al., 2011; Klimesch, 2012; Klimesch, Sauseng & Hanslmayr, 2007) that alpha oscillations gate the flow of information in corticothalamic and intra-cortical circuits, and in turn, shape attention and perception.

Uusberg et al. (2013) tested this hypothesis by measuring alpha activity while subjects viewed emotionally valenced stimuli, and reported increased alpha-band event-related desynchronisation (ERD) in response to aversive compared to neutral stimuli. Analogous research corroborates this finding, suggesting alpha ERD to emotional information reflects enhanced neuronal excitation induced by affective attention, whereby attention is focused on emotional stimuli for heightened sensory processing (Aftanas et al., 2002; Güntekin & Başar, 2007; Onoda et al., 2007). This interpretation is consistent with behavioural findings of valence-induced changes in attention at encoding and superior performance for emotional stimuli at recall (Riggs, McQuiggan, Farb, Anderson & Ryan, 2011).

### 1.2. Attention in sleep-related emotional memory consolidation

Evidence suggests typical nocturnal sleep or daytime naps facilitate memory formation compared to wakefulness (Born & Wilhelm, 2012; Mednick Nakayama & Stickgold, 2003; Wagner, Gais & Born, 2001), and particularly for emotional memories (Diekelmann et al., 2009; Groch et al., 2013; Hutchison & Rathore, 2015; Payne et al., 2015). Of the physiological processes active during sleep, rapid eye movement sleep (REM) has been implicated in emotional memory consolidation (Groch et al., 2013; Wagner et al., 2001). This enhancement is thought to be facilitated by REM theta oscillations which represent homeostatic processes of emotional brain regulation and have been linked to the synchronisation of emotional information between the amygdala and hippocampus, regions involved in emotional long-term memory (Diekelmann et al., 2009; Hutchison & Rathore, 2015; Prehn-Kristensen et al., 2013). However, recent findings suggest REM neurophysiology does not solely explain emotional memory consolidation (Morgenthaler et al., 2014). Alternatively, slow-wave sleep (SWS) is thought to facilitate the consolidation of emotional information through decreasing synaptic connectivity, ultimately pruning unnecessary neural connections and thus refining memory representations (Morgenthaler et al., 2014; Payne et al., 2015; Walker, 2010). As such, it is possible that both SWS oscillations and REM theta oscillations are important for the consolidation of emotional memory (Cunningham, Chambers & Payne, 2014; Hutchison & Rathore, 2015).

This is supported by emerging evidence that indicates shorter sleep durations, such as an afternoon nap, are equally effective in stabilising new information as nocturnal sleep paradigms (Nishida, Pearsall, Buckner & Walker, 2009; Payne et al., 2015). An afternoon nap occurs at a different circadian phase than nocturnal sleep and is typically dominated by NREM sleep (Payne et al., 2015). It also allows relatively better control of potential time-of-day effects on performance, and if sleep facilitates long-term memory through a process of active consolidation then, emotional memory should benefit from an afternoon nap compared to an equivalent period of wake.

Studies investigating the relationship between sleep and emotional memory consolidation have not accounted for attention during encoding, limiting the ability to establish links between attention-related memory enhancement and sleep-related memory consolidation. This limitation is emphasised by Saletin & Walker (2012), who argue that sleep-dependent memory consolidation involves a discriminatory mechanism that is determined by salient cues at encoding, such as emotionality, resulting in a prioritisation of emotional over neutral information. As attentional biases toward emotional information lead to enhanced memory performance (MacKay et al., 2004; Riggs et al., 2011), alpha desynchronisation at encoding may serve to modulate emotional memory consolidation (Everaert & Koster, 2015; Smith et al., 2004; Wang & Bastiaansen, 2014). Further, if sleep plays an active role in memory consolidation, and if alpha rhythms reflect modulations in attention and perception, alpha oscillatory activity at encoding should further modulate memory performance after a period of sleep compared to an equivalent period of wake.

### 1.3. Current Study

The present study examined whether attention modulates memory consolidation for emotional information across a period of sleep versus wakefulness. Specifically, this study aimed to determine whether individual alpha oscillatory activity at encoding facilitates sleep-related memory consolidation and whether this facilitation is higher for emotional compared to neutral stimuli.

It was hypothesised that condition (sleep, wake), attention (alpha spectral power) and emotional valence (positive, negative, neutral) would interact in their effect on emotional memory performance. Specifically, it was predicted that: (1) memory performance would be greater after sleep compared to wake; (2) memory performance would be greater for emotional compared to neutral information and this effect would be accentuated after sleep compared to wake; (3) greater attention, as indexed by alpha oscillatory activity, would be associated with greater memory performance, and; (4) differences in memory performance relative to condition and emotional valence would be modulated by alpha activity.

## 2. Method

### 2.1. Participants

Participants included 22 right-handed healthy adults (10 male) ranging from 18 – 41 years old (mean age = 27.3). A power analyses using G*Power 3 (Faul, Erdfelder, Lang & Buchner, 2007) of a previous study examining the impact of sleep on emotional memory (partial η^2^ = .20 based on Payne et al., 2008) suggested a sample size of 12 would be adequate to detect similar sized effects in a repeated measures design (*β*=80, *α*=.05). All participants reported normal or corrected-to-normal vision and hearing and had no current or past psychiatric conditions, substance dependence or abuse, intellectual impairment and were not taking medication that influenced sleep and neuropsychological measures. All participants provided informed consent and received a $40 honorarium. One participant did not return during the second condition, resulting in a final sample size of 21 (9 males; mean age = 26; age range = 18 – 41). Ethics for this study was granted by the University of South Australia’s Human Research Ethics committee (I.D: 0000032556).

### 2.2. Design

This study was a repeated measures within-subjects experimental design with two conditions (sleep, wake). Each condition was counterbalanced across participants and separated by one week to control for condition order effects and to avoid interference between task sets. Conditions included:

a. Sleep condition: Participants underwent learning with an immediate retrieval task followed by a 2hr sleep opportunity. A delayed retrieval task occurred thirty minutes after waking.
b. Wake condition: Participants underwent learning with an immediate retrieval task. This was followed by a delayed retrieval task after a 2hr wake period.

### 2.3. Materials and measures

#### 2.3.1. 2.3.1.Demographic measures

Participants completed a paper questionnaire containing questions on age, sex, ethnicity, highest level of education achieved and recent (<24hr) alcohol and caffeine consumption, as caffeine and alcohol are known to influence performance on cognitive tasks (Keenan, Tiplady, Priestley & Rogers, 2014).

#### 2.3.2. Screening and control measures

The Pittsburgh Sleep Quality Index (PSQI; Buysse, Reynolds, Monk, Berman & Kupfer, 1989) was used to screen for sleep quality. Participants’ PSQI scores ranged from 1 – 5 (*M* = 3.4, *SD* = 1.50), indicating good sleep quality. The WASI-II was used to ensure average intellectual ability, as intelligence may influence memory retention and performance on memory tasks (Conway, Kane & Engle, 2003). The WASI-II provides an estimate of full-scale IQ (FSIQ). Participants’ mean FSQI score was 114 (*SD* = 16.44), placing participants in a high range of intellectual functioning (Wechsler, 2011).

#### 2.3.3. Visual analogue scale for sleepiness

The visual analogue scale for sleepiness (VASS) was used to control for sleepiness. The VASS is comprised of a 100mm visual analogue scale with “sleepy/drowsy” and “alert/awake” for endpoints to denote a continuum of state sleepiness. Participants indicated where on the line they judged their current state of sleepiness prior to the start and end of each learning and recall session. Scores were determined as mm distance from the left pole to the participants’ mark, indicating degree of sleepiness as a percentage, with lower scores indicating greater sleepiness.

#### 2.3.4. Polysomnography (PSG)

PSG was recorded using the Compumedics Grael High-Definition PSG 24-bit amplifier (Compumedics Pty Ltd., Melbourne, Australia). Electrodes were arranged according to the International 10-20 System (American Electroencephalographic Society, 1994) at the following locations: FP1, FP2, F3, F4, C3, C4, T7, T8, P3, P4, P7, P8, O1, O2. In addition to left and right electro-oculography (EOG), sub-mental electromyography (EMG) and electrocardiography (ECG), EEG was recorded and referenced to contralateral mastoids and sampled at a rate of 512 Hz with a bandpass filter from DC to 143 Hz. All impedances were kept at or below 10kΩ throughout recording periods. All sleep data were scored by an experienced sleep technician according to standardised criteria (Berry et al., 2012) with EEG viewed with a high pass filter of 0.3 Hz and a low pass filter of 35 Hz. The following sleep parameters were derived from PSG recordings: time in bed, total sleep time (TST), sleep onset latency (SOL; time from lights out to the first epoch of sleep), REM onset latency, sleep efficiency (SE) [(total sleep time/time in bed) x 100], wake after sleep onset, total arousal index, duration and percent of TST spent in each sleep stage.

### 2.4. EEG paradigm (attentional state)

EEG was continuously recorded during a visual Conjunctive Continuous Performance Task (CCPT-V), which was used to elicit sustained attention derived from target discrimination in the visual modality. The CCPT-V began with 15 practice trials followed by 320 trials and took approximately eight minutes to complete (Shalev, Ben-Simon, Mevorach, Cohen & Tsal, 2011). Stimuli were coloured geometric shapes and were presented in the centre of a computer monitor. Target probability (red square) was 30%, non-red square probability was 17.5%, red non-square probability was 17.5%, and all other shape (neither a square nor red) probability was 35%. Each inter-stimulus interval (ISI) appeared in 25% of trials and both the stimuli and ISIs were randomly selected (Shalev et al., 2011). Participants were instructed to press a key on a standard keyboard whenever a target appeared and to refrain from responding to all other stimuli. An example stimulus train is presented in Figure 1.

**Figure 1.**
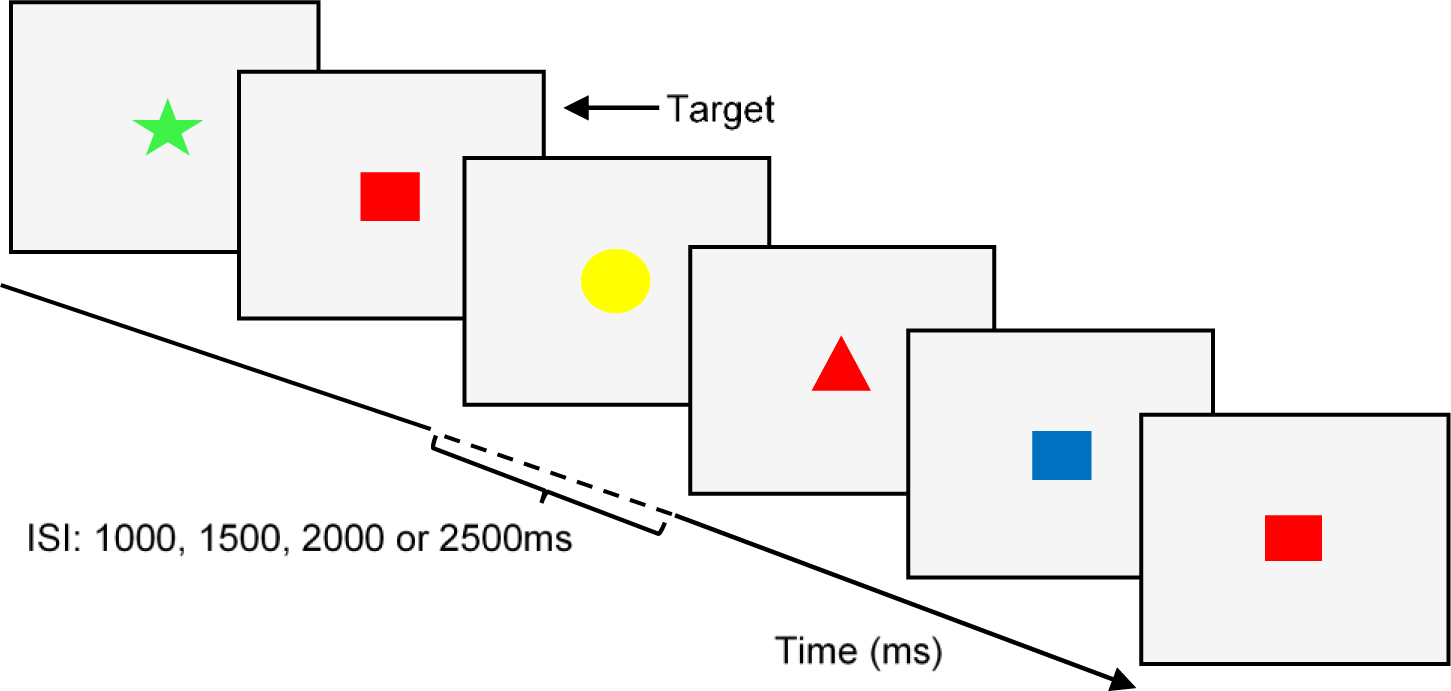
Example of a sequence of trials on the Conjunctive Continuous Performance Task – Visual. Stimulus presentation was 100ms, separated by an ISI of 1000, 1500, 2000 or 2500ms.

### 2.5. Memory task

480 pictures were used from the International Affective Picture System (IAPS; Lang, Bradley & Cuthbert, 2008), which were divided into two parallel sets (Set A, Set B), counterbalanced across the sleep and wake conditions. The IAPS are rated for emotionality based on two dimensions (valence: 1=unpleasant, 9=pleasant; arousal: 1=calm, 9=excited) by a normative adult sample (Lang et al., 2008). Mean valence and arousal ratings of the stimuli have high internal consistency (*a*=.94) and split-half reliability (*r*s=.94).

Stimuli were displayed via OpenSesame Software version 2.9.7 (Mathôt, Schreij & Theeuwes, 2012) and were categorised into three groups (i.e., negative, neutral, positive) according to their mean valence values. Negative pictures had a mean valence rating of 3.27 (*SD* = 0.03), neutral pictures had a mean valence rating of 5.02 (*SD* = 0.03), and positive pictures had a mean valence rating of 7.00 (*SD* = 0.03). All stimuli were counterbalanced based on valence and arousal.

During the learning tasks, for each set, 120 stimuli, each containing 40 neutral, 40 positive and 40 negatively valenced images, were presented sequentially for participants to learn, followed by an immediate retrieval task to gain a recognition baseline. A delayed retrieval task occurred post experimental conditions. The immediate and delayed retrieval tasks contained 120 previously seen pictures intermixed with 120 new (distractor) pictures (see Figure 2 for schematic representation). Following presentation, participants judged whether pictures were old or new. Pictures were pseudo-randomised at each time of testing, such that no more than two pictures of the same emotion followed. Testing time for each experimental task was approximately 20 – 30 minutes.

**Figure 2.**
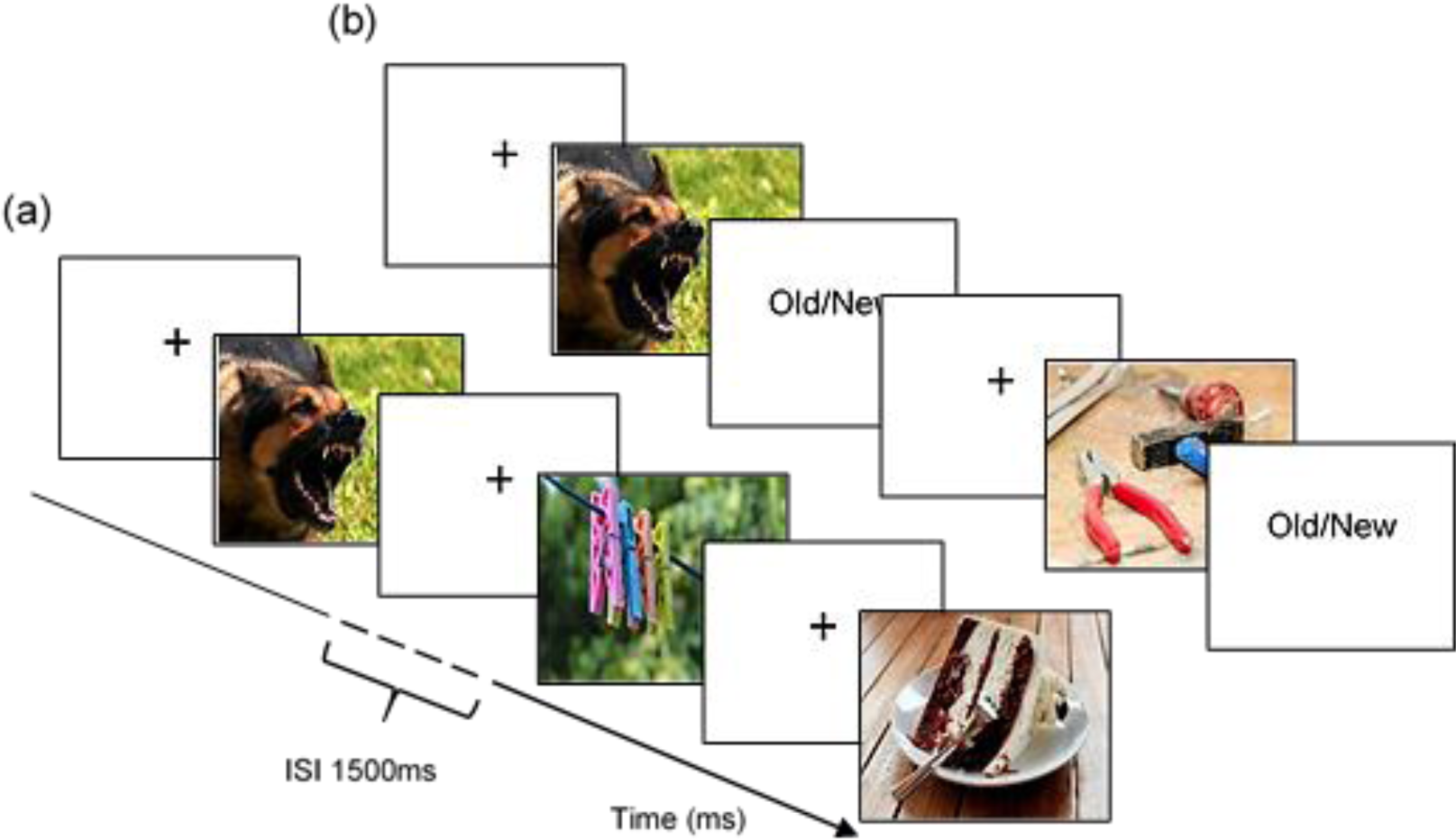
(a) Schematic representation of the learning task. Stimuli were presented for 1000ms to allow for conscious processing whilst maintaining an appropriate degree of difficulty. Each stimulus was preceded by a 500ms fixation cross with a temporal jitter of ± 100ms and with an ISI of 1500ms; (b) Schematic representation of the retrieval task. Stimuli were presented until button press with a timeout at 5000ms, preceded by a 500ms fixation cross with a jitter of ± 100ms. Participants then indicated whether the stimulus was old or new. Images are taken from Creative Commons for illustrative purposes, as the IAPS are restrictively licensed and not available for general distribution.

### 2.6. Procedure

Participants came to the laboratory at approximately 11:45hr, and were taken through to the testing rooms for PSG/EEG set-up. Baseline EEG activity was recorded during quiet sitting with eyes open (focussing on a fixation cross centred on a computer monitor) and eyes closed for two minutes, respectively. Participants then completed the CCPT-V and learning task by approximately 13:30hr, followed by the immediate recall task.

During the wake condition, participants remained in the laboratory and were administered the WASI-II. During the sleep condition, participants were given a 120-minute sleep opportunity between the hours of 14:30hr and 16.30hr. Nap success rate was 100%. PSG data was used to verify sleep periods. Participants engaged in non-strenuous activity for 30 minutes after the nap until testing to alleviate inertia effects on memory performance (Cairney, Durrant, Hulleman & Lewis, 2014; Payne et al., 2008; Tassi & Muzet, 2000). At approximately 17:00hr, participants completed the delayed retrieval task (see Figure 3 for study protocol).

**Figure 3.**
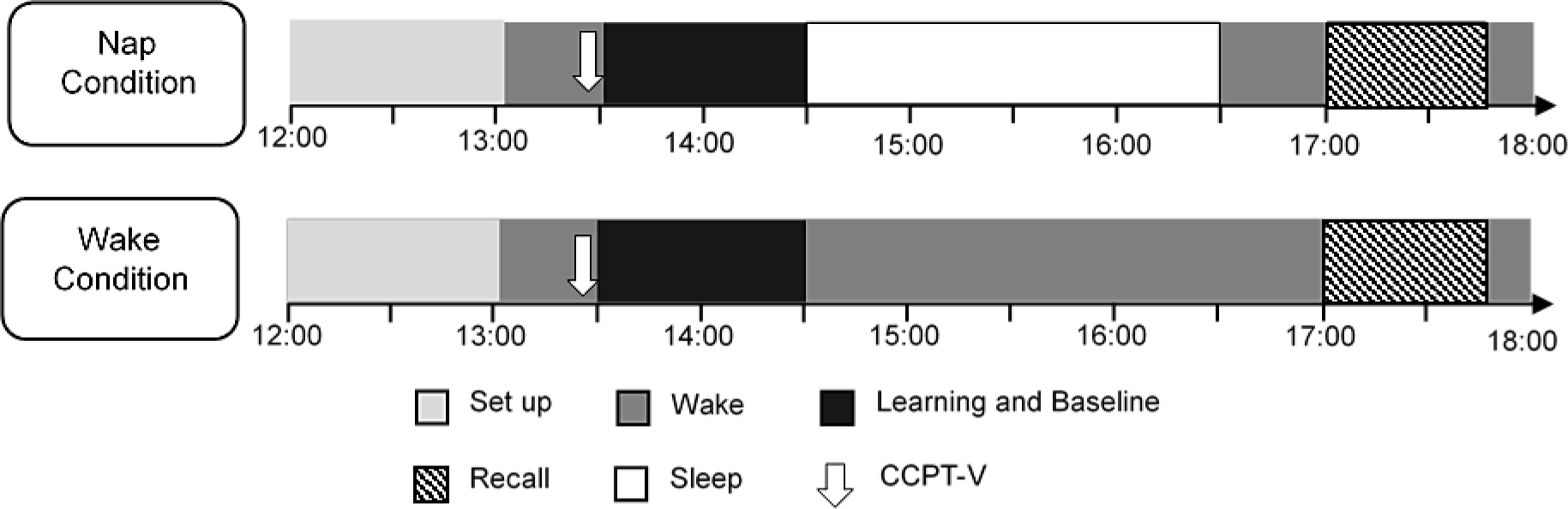
Diagram representing the time course of the experimental conditions (sleep, wake) and testing session (learning, retrieval). Timing of the CCPT-V is shown in downward arrows at approximately 13.25hr.

## 3. Data Analysis

### 3.1. Behavioural data

The mean response time (ms) to correct responses (M-RT), the percentage of commission errors, and percentage of correct responses were determined from CCPT-V performance. M-RT reflects response time, while commissions and the percentage of correct responses reflect aspects of the accuracy of responses (Shalev et al., 2011). Memory performance was calculated on a trial-by-trial basis, determined by whether participants correctly identified target and distractor stimuli.

### 3.2. EEG pre-processing and spectral analysis

All EEG analysis procedures were conducted using customised scripts programmed in MATLAB^®^ (R2015a, The MathWorks, Inc., Natick, MA, USA). EOG and parieto-occipital EEG channels (P3-M2, P4-M1, O1-M2, and O2-M1) were imported into MATLAB via the EEGLAB toolbox (v13.6.5b; Delorme & Makeig, 2004) and subjected to zero-phase, finite impulse response highpass (passband edge: 1 Hz, -6 dB cutoff: 0.5 Hz) and lowpass (passband edge: 40 Hz, -6 dB cutoff: 45 Hz) Hamming-windowed sinc filters (implemented via the pop_eegfiltnew function of the firfilt plugin; v1.6.1). Datasets were then divided into two separate segments corresponding to the eyes-closed resting-state and CCPT-V portions of the EEG recording. An automated artifact detection routine was applied to both segments in order to exclude blinks and other sources of signal contamination. The peak-to-peak threshold for artifact rejection was set at +/− 75 μV, and applied within a 500ms sliding window (50% overlap). EOG channels were removed from the data following artifact rejection.

One-sided PSD estimates were generated via the MATLAB implementation of Welch’s modified periodogram method (Welch, 1967). This technique divides the signal into equal segments, multiplies each segment by a tapering window of identical length, computes the fast Fourier transform of this product, and averages across the resulting periodograms to estimate the PSD. We applied a 2048 sample (4s) Hamming window with 50% overlap across segments, yielding a frequency resolution of 0.244 Hz. For each channel, PSD estimates were normalised by dividing each power estimate within the passband by the average (mean) power of the spectrum. This step was designed to facilitate comparison of spectral activity across conditions.

Channel-wise PSD estimates derived over the duration of the CCPT-V task were averaged for each participant, and summed across alpha-band frequency bins to summarise the quantity of alpha-band activity per condition. Alpha bandwidth was defined as 2 Hz above and below the participant’s individual alpha frequency (IAF). Two indices of IAF (peak frequency and centre of gravity) were calculated from eyes-closed resting-state recordings (estimates from the sleep and wake conditions were grand-averaged, unless only one estimate was available).

IAF estimates were obtained using *restingIAF* v1.0, an open-source package available from https://github.com/corcorana/restingIAF. This automated IAF estimation routine uses a Savitzky-Golay filter (frame length = 11 frequency bins, polynomial degree = 5) to smooth the PSD. It then searches the first derivative of the smoothed PSD for evidence of peak activity within a defined frequency interval (here, 7–13 Hz), and parameterises the bounds of identified peak components to compute the centre of gravity. Given the relatively low number of parieto-occipital channels available for analysis (i.e. 4), the minimum number of valid channel estimates required to estimate IAF was set to 1. All other analysis parameters were as per default settings (see Corcoran, Alday, Schlesewsky, & Bornkessel-Schlesewsky, 2018).

### 3.3. Statistical analysis

Data were entered into *R* version 3.4.0 (*R* Core Team, 2017) for Windows, and the *lme4* package for linear mixed-effects models (Bates, Maechler, Bolker & Walker, 2015). A generalised linear mixed effects model (GLMM) with a logit link function fit by maximum likelihood was used to examine the relationship between alpha oscillatory activity, emotional valence and sleep on memory consolidation.

Mixed models are an appropriate method for analysing data from repeated measure designs, as these designs are grouped by subject and appropriately account for within and between subject variance (Judd, Westfall & Kenny, 2012; Van Dongen, Olofsen, Dinges & Maislin, 2004). Further, logit mixed models appropriately account for binomial response variables (Jaeger, 2008). As such, a logit mixed model is particularly suited to the current data due to: (a) large inter-individual differences in IAF estimates; (b) inter-individual differences in response to the emotional valence of each stimulus, and; (c) the use of categorical response variables (i.e., binomial old/new memory responses) as a measure of memory performance.

Condition (sleep, wake), Valence (positive, negative, neutral), Time (baseline, delayed) and Alpha were specified as fixed effects. Alpha power was centred and scaled via a z-transformation prior to inclusion in the GLMM. The model included random-effects terms for items and participant on the intercept. Memory performance (correct-by-trial) was specified as the dependent variable. Akaike Information Criterion (AIC; Akaike, 1974), Bayesian Information Criterion (BIC; Schwarz, 1978) and log-likelihood were used to assess the fit of the GLMM. All models were fully saturated with main and interaction effects for Condition, Valence, Time and Alpha, while Type II Wald Chi-Square (χ2) tests from the *car* package in *R* (Fox & Weisberg, 2011) were used to provide *p*-value estimates for each of the factors. Pearson correlations were computed between the percentage of time spent in N2, SWS and REM on memory performance to investigate the relationship between sleep and emotional memory consolidation. All *p*-values are 2-tailed, with statistical significance determined at α = 0.05. All data are presented as mean and standard deviation (SD) unless indicated otherwise, and effects were plotted using the package *effects* (Fox, 2003) and ggplot2 (Wickham, 2009).

## 4. Results

### 4.1. Preliminary analyses

Preliminary analyses were conducted to determine: (1) if there were any differences in behavioural performance on the CCPT-V measures between conditions; (2) if there were any differences in alpha power between the two conditions; (3) whether there was a difference in levels of sleepiness between the two conditions prior to the baseline and delayed recall tasks, and; (4) the sleep characteristics of the nap. Since the IAF estimators were highly correlated (r = .96, *p* < .001, 95% CI = [.90, .98]), all reported alpha power estimates were derived from individual bandwidths centred on the peak alpha frequency (see supplementary materials for reanalysis of power estimates in relation to the alpha centre of gravity). Peak alpha frequency estimates from both conditions are displayed in Figure 4.

**Figure 4.**
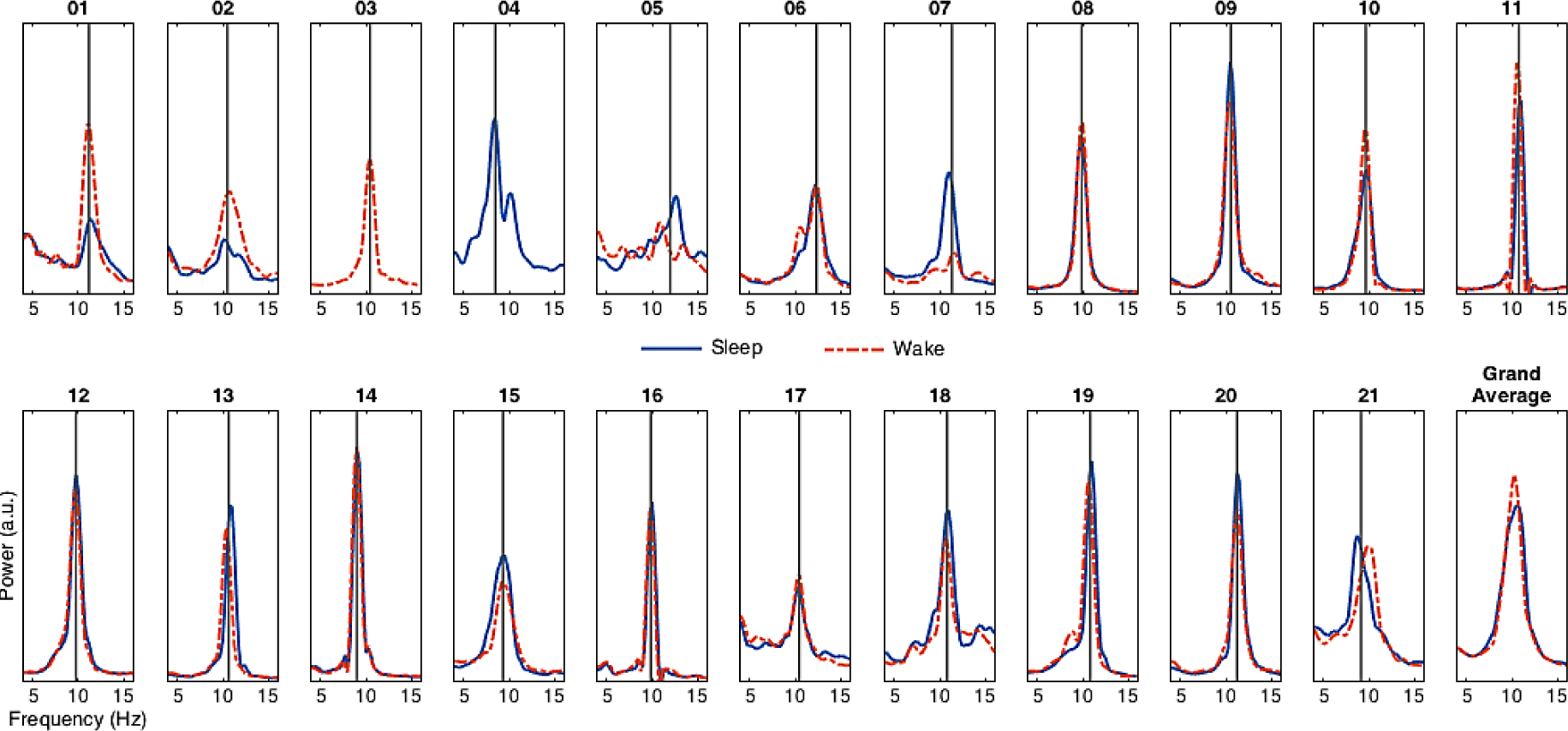
Individual (peak) alpha frequency (IAF) estimates from eyes-closed resting-state recordings prior to the Sleep (blue, solid lines) and Wake (red, broken lines) conditions. Vertical line indicates the mean IAF estimate used to locate the individualised alpha bandwidth for the main analysis. Note that resting-state data were missing for participant 4 during the Wake condition. Participant 3 did not demonstrate evidence of a distinct alpha peak during the Sleep condition resting-state recording, hence IAF was estimated on the basis of the Wake condition peak frequency. a.u.: arbitrary units.

Paired sample *t*-tests conducted between the three CCPT-V performance measures demonstrated no significant difference between conditions in reaction times (*t*(20)=.36, *p*=.72), percentage of commissions (*t*(20)=.60, *p*=.60) and percentage of correct responses (*t*(20)=.67, *p*=.50). See Table 1 for means and standard deviations of the CCPT-V performance measures. Similarly, paired samples t-test indicated relative alpha power did not differ between the sleep (*M*=30.81, *SD*=17) and wake (*M*=28.31, *SD*=16.16) conditions (*t*(20)=.92, *p*=.36).

**Table 1.** Means and Standard Deviations of Reaction Times, the Percentage of Commissions and the Percentage of Correct Responses for the CCPT-V between the Sleep and Wake Conditions.

| CCPT-V Variables | Condition |  |
| --- | --- | --- |
|  | Sleep M (SD) | Wake M (SD) |
| Reaction Time (ms) | 422.35 (39.11) | 419.14 (41.92) |
| % of Commissions | 0.74 (0.51) | 0.81 (0.65) |
| % of Correct Responses | 99.55 (1.02) | 99.21 (2.02) |

Subjects were significantly sleepier during the Sleep (*M*=54.57, *SD*=18.53) than the Wake (*M*=64.43, *SD*=21.23) condition prior to the learning and baseline recall tasks (*t*(20 = 2.26, *p*<.001, *d* =-.46). There was no difference between the conditions at delayed recall (Sleep: *M*=61.47, *SD*=16.42, Wake: *M*=67.64, *SD*=15, *t*(20 = -1.23, p>.05, *d* =-.39). The CCPT-V data also showed no significant difference, suggesting attentional state was equivalent between the two conditions at delayed recall (*t*(20 = -1.65, *p*>.05, *d* = -.19). A linear regression was conducted to determine whether the greater levels of sleepiness in the Sleep group impacted baseline memory performance. The results of the regression indicate that there was no significant effect of self-reported sleepiness on baseline memory performance (*β* = *-0.71*, p = .66, *R*^2^ = -0.04).

Sleep variables of total sleep time, sleep onset latency, wake after sleep onset and the amount of time and percentage of sleep spent in stage 1 (N1), stage 2 (N2), SWS and REM are reported in Table 2. Sleep data show the expected proportion of NREM sleep stages (i.e., N1, N2 and SWS) and minimal REM sleep (Payne et al., 2015). That is, although 81 percent of participants experienced REM, average time spent in this stage was only 10 minutes (*SD* = 7.71).

**Table 2.** Means, Standard Errors and Ranges for Sleep Parameters.

| Sleep Parameters | Minutes |  |
| --- | --- | --- |
|  | Mean (SEM) | Range |
| TST (min) | 91.45 (5.76) | 23 – 119 |
| SOL (min) | 13.64 (3.07) | 1 – 53.5 |
| WASO (min) | 15.10 (3.86) | 2.5 – 65.5 |
| N1 (%TST) | 8.12 (1.22) | 0.4 – 19.2 |
| N2 (%TST) | 51.78 (2.99) | 28.7 – 87 |
| SWS (%TST) | 31.33 (3.60) | 0 – 55.4 |
| REM (%TST) | 8.77 (1.76) | 0 – 32 |

*Note.* SEM = standard error of the mean. TST = total sleep time; SOL = sleep onset latency; WASO = wake after sleep onset; N1 = stage 1; N2 = stage 2; SWS = slow wave sleep; REM = rapid eye movement sleep; min = minutes.

### 4.2. Primary analysis

All significant effects and interactions of the GLMM of memory performance of correct-by-trial are given in Table 3. GLMM results indicate a significant effect of Condition, with factor level contrasts revealing memory performance was significantly greater after wake compared to sleep. There was no effect of Valence, translating into a statistically non-significant difference in memory performance between negative, neutral and positive stimuli. There was a significant effect of Time, with memory performance significantly greater at Baseline compared to Delayed testing (see Figure 5). Finally, a significant effect of Alpha was indicated for memory performance, such that enhanced memory performance was predicted by increased alpha power prior to encoding.

**Figure 5.**
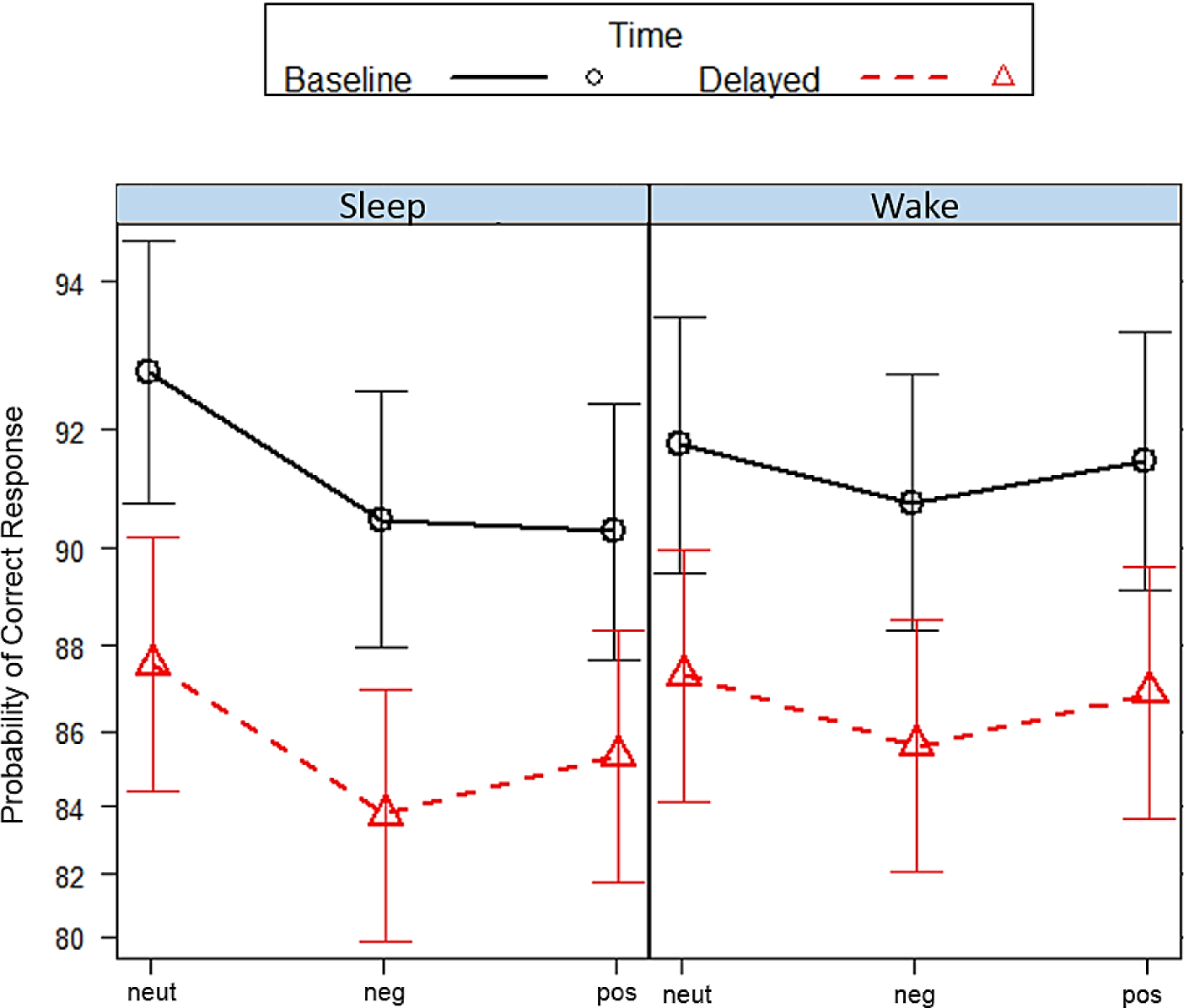
Plot of effects for Condition (Sleep, Wake) on memory for neutral, negative and positively valenced stimuli at baseline and delayed testing. Bars indicated 83% confidence intervals.

**Table 3.**
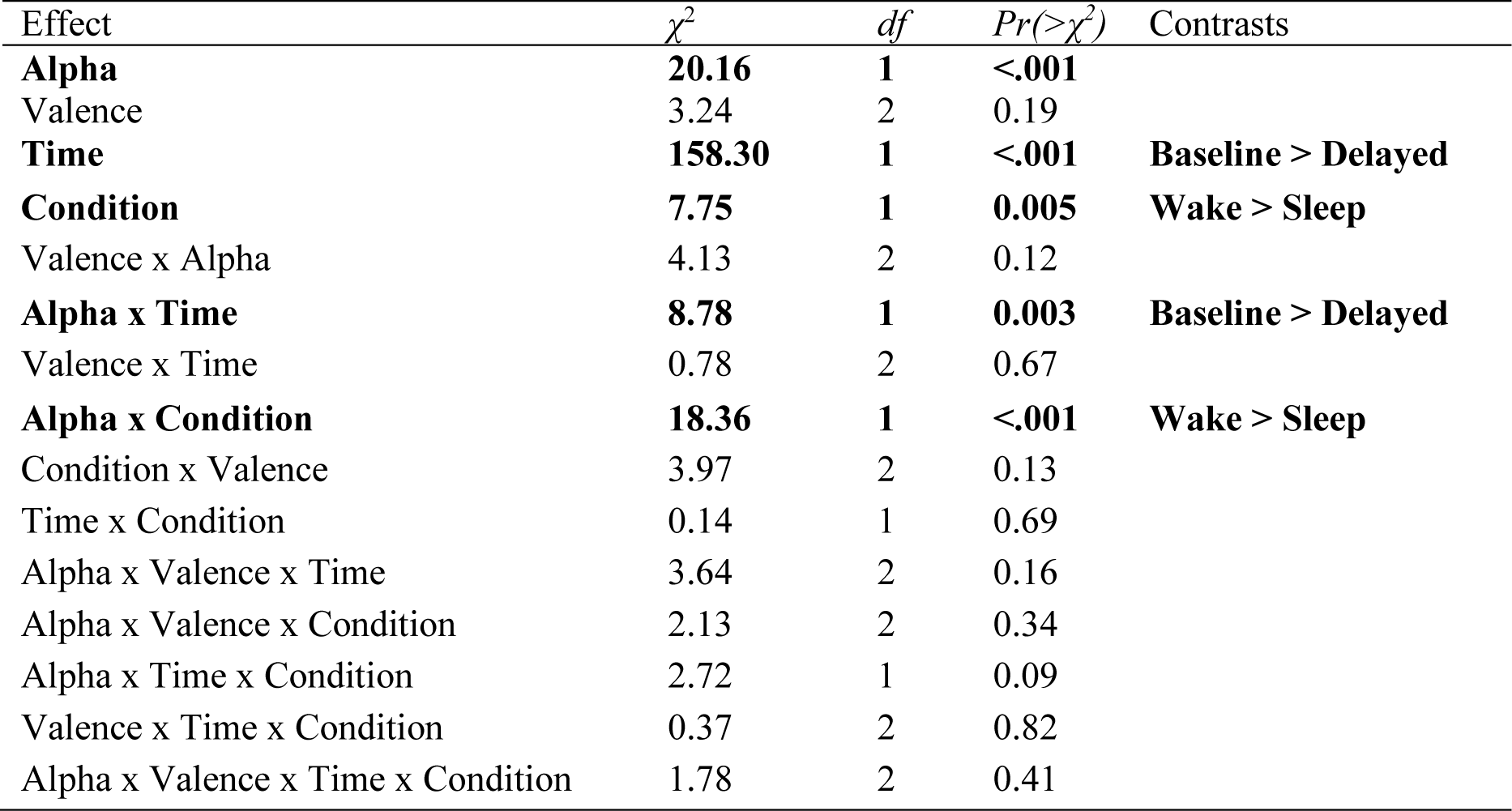
Type-II Wald Tests for all Main Effects and Interactions with Chi-Square ratio values, Degrees of Freedom and Statistical Significance. Final Column Display Significant Factor Level Contrasts. Significant effects and interactions are bold-faced.

| Effect | $\chi^2$ | df | $Pr(>\chi^2)$ | Contrasts |
| --- | --- | --- | --- | --- |
| **Alpha** | **20.16** | **1** | **<.001** |  |
| Valence | 3.24 | 2 | 0.19 |  |
| **Time** | **158.30** | **1** | **<.001** | **Baseline > Delayed** |
| **Condition** | **7.75** | **1** | **0.005** | **Wake > Sleep** |
| Valence x Alpha | 4.13 | 2 | 0.12 |  |
| **Alpha x Time** | **8.78** | **1** | **0.003** | **Baseline > Delayed** |
| Valence x Time | 0.78 | 2 | 0.67 |  |
| **Alpha x Condition** | **18.36** | **1** | **<.001** | **Wake > Sleep** |
| Condition x Valence | 3.97 | 2 | 0.13 |  |
| Time x Condition | 0.14 | 1 | 0.69 |  |
| Alpha x Valence x Time | 3.64 | 2 | 0.16 |  |
| Alpha x Valence x Condition | 2.13 | 2 | 0.34 |  |
| Alpha x Time x Condition | 2.72 | 1 | 0.09 |  |
| Valence x Time x Condition | 0.37 | 2 | 0.82 |  |
| Alpha x Valence x Time x Condition | 1.78 | 2 | 0.41 |  |

There was a significant Alpha x Condition interaction. Factor level contrasts revealed Alpha modulated memory performance greater in the wake compared to the sleep condition, such that memory performance after a period of wake was enhanced with increased alpha power prior to encoding (see Figure 6). The Condition x Valence and Valence x Time interactions were nonsignificant, indicating memory for negative, neutral and positive stimuli did not differ between the sleep and wake conditions at baseline and delayed testing, respectively. Similarly, the Valence x Alpha interaction was nonsignificant, suggesting enhanced baseline attention did not modulate memory for negative, neutral and positive stimuli. The Alpha x Time interaction was significant, indicating that increased alpha power prior to encoding was a stronger predictor of baseline relative to delayed memory performance; however, alpha power prior to encoding remained a strong predictor of memory performance at Delayed recall, as illustrated in panel B of Figure 6. By contrast, the Time x Condition interaction was nonsignificant, indicating that the change in memory performance from baseline to delayed testing did not differ between the Sleep and Wake conditions. All three and four way interactions were nonsignificant.

**Figure 6.**
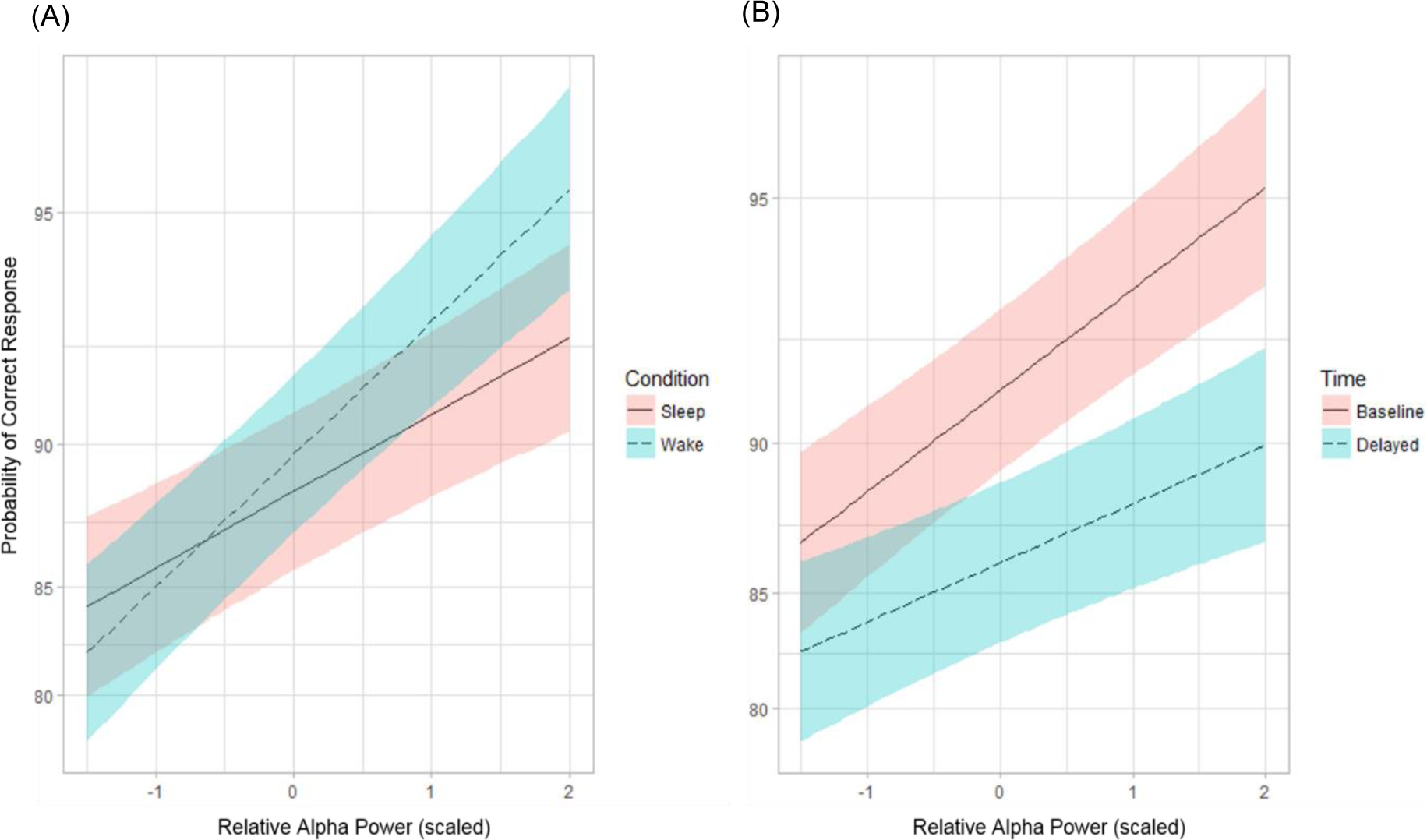
(A) Interaction between relative alpha power and the sleep (red) and wake (blue) conditions on memory performance at delayed recall. (B) Interaction between relative alpha power on memory performance at baseline (red) and delayed (blue) testing. Higher values on the y-axis indicate greater memory performance, while higher values on the x-axis indicate greater alpha synchronisation over left (O1-P3) and right (O2-P4) occipital-parietal electrodes during the CCPT-V prior to the learning task. The blue and red shaded space indicate the 83% confidence interval.

**Table 4.**
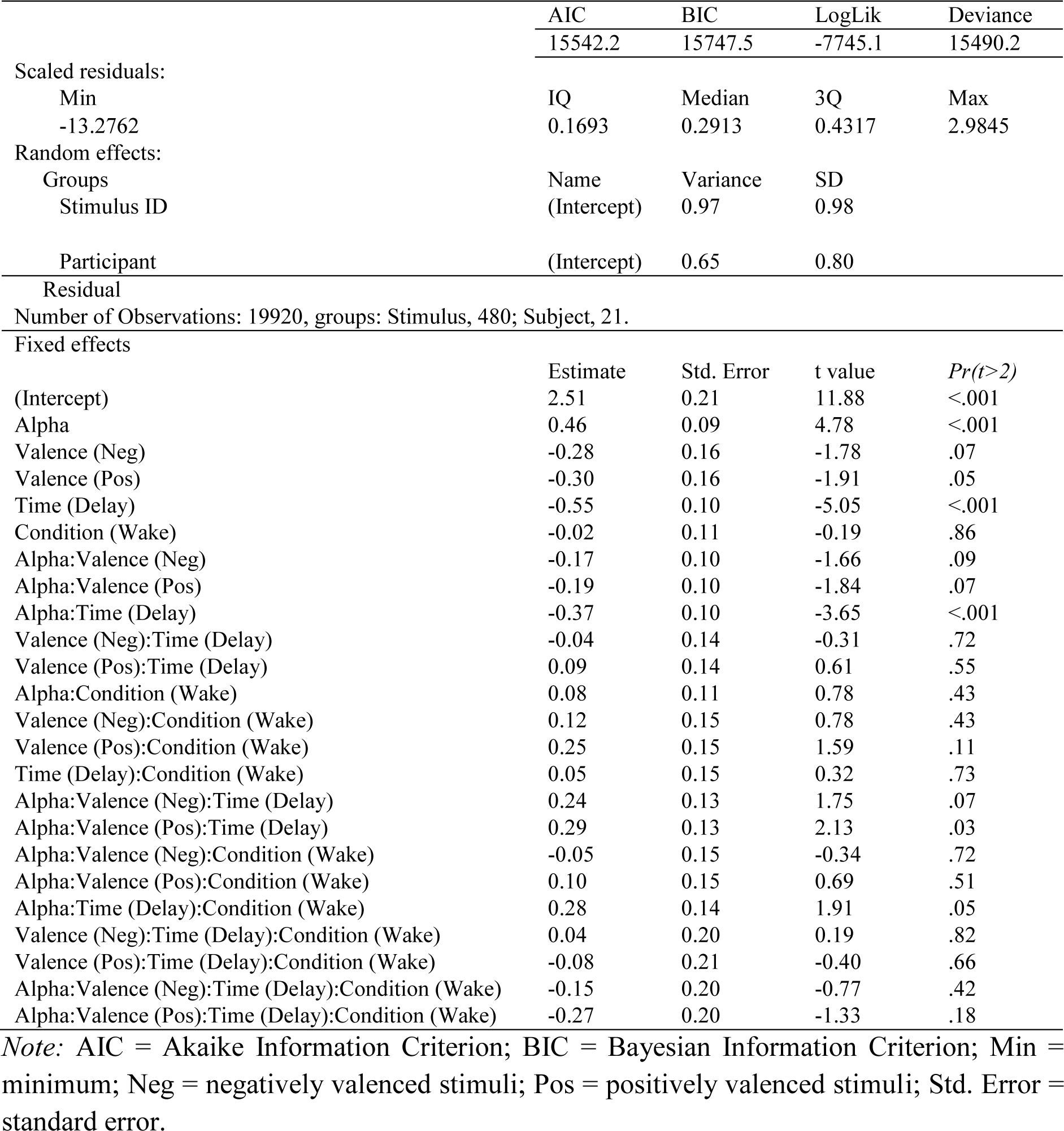
Generalised Linear Mixed Model Fit by Maximum Likelihood (Laplace Approximation) with Alpha as a Neurophysiological Measure of Attentional State Prior to Encoding.

|  | AIC | BIC | LogLik | Deviance |
| --- | --- | --- | --- | --- |
|  | 15542.2 | 15747.5 | -7745.1 | 15490.2 |
| Scaled residuals: |  |  |  |  |
| Min | IQ | Median | 3Q | Max |
| -13.2762 | 0.1693 | 0.2913 | 0.4317 | 2.9845 |
| Random effects: |  |  |  |  |
| Groups | Name | Variance | SD |  |
| Stimulus ID | (Intercept) | 0.97 | 0.98 |  |
| Participant | (Intercept) | 0.65 | 0.80 |  |
| Residual |  |  |  |  |
| Number of Observations: 19920, groups: Stimulus, 480; Subject, 21. |  |  |  |  |
| Fixed effects | Estimate | Std. Error | t value | *Pr*(*t*>2) |
| (Intercept) | 2.51 | 0.21 | 11.88 | <.001 |
| Alpha | 0.46 | 0.09 | 4.78 | <.001 |
| Valence (Neg) | -0.28 | 0.16 | -1.78 | .07 |
| Valence (Pos) | -0.30 | 0.16 | -1.91 | .05 |
| Time (Delay) | -0.55 | 0.10 | -5.05 | <.001 |
| Condition (Wake) | -0.02 | 0.11 | -0.19 | .86 |
| Alpha:Valence (Neg) | -0.17 | 0.10 | -1.66 | .09 |
| Alpha:Valence (Pos) | -0.19 | 0.10 | -1.84 | .07 |
| Alpha:Time (Delay) | -0.37 | 0.10 | -3.65 | <.001 |
| Valence (Neg):Time (Delay) | -0.04 | 0.14 | -0.31 | .72 |
| Valence (Pos):Time (Delay) | 0.09 | 0.14 | 0.61 | .55 |
| Alpha:Condition (Wake) | 0.08 | 0.11 | 0.78 | .43 |
| Valence (Neg):Condition (Wake) | 0.12 | 0.15 | 0.78 | .43 |
| Valence (Pos):Condition (Wake) | 0.25 | 0.15 | 1.59 | .11 |
| Time (Delay):Condition (Wake) | 0.05 | 0.15 | 0.32 | .73 |
| Alpha:Valence (Neg):Time (Delay) | 0.24 | 0.13 | 1.75 | .07 |
| Alpha:Valence (Pos):Time (Delay) | 0.29 | 0.13 | 2.13 | .03 |
| Alpha:Valence (Neg):Condition (Wake) | -0.05 | 0.15 | -0.34 | .72 |
| Alpha:Valence (Pos):Condition (Wake) | 0.10 | 0.15 | 0.69 | .51 |
| Alpha:Time (Delay):Condition (Wake) | 0.28 | 0.14 | 1.91 | .05 |
| Valence (Neg):Time (Delay):Condition (Wake) | 0.04 | 0.20 | 0.19 | .82 |
| Valence (Pos):Time (Delay):Condition (Wake) | -0.08 | 0.21 | -0.40 | .66 |
| Alpha:Valence (Neg):Time (Delay):Condition (Wake) | -0.15 | 0.20 | -0.77 | .42 |
| Alpha:Valence (Pos):Time (Delay):Condition (Wake) | -0.27 | 0.20 | -1.33 | .18 |

*Note:* AIC = Akaike Information Criterion; BIC = Bayesian Information Criterion; Min = minimum; Neg = negatively valenced stimuli; Pos = positively valenced stimuli; Std. Error = standard error.

#### 4.2.1. Relationship between sleep stages and memory performance

Pearson correlations were conducted to assess the relationship between sleep and memory performance (see Table 5 for a summary of the correlation coefficients). None of the relationships between the percentage of time spent in N2, SWS and REM on the consolidation of negative, neutral and positive stimuli were significant. Similarly, the relationship between the percentage of time spent in these sleep stages on overall memory performance (i.e. collapsed across positive, negative and neutral stimuli) was not significant. However, although nonsignificant, there was a small-to-moderate relationship between the percent of time spent in N2 and memory for emotional (positive and negative) stimuli. See Figure 7 for scatterplots illustrating the relationship between sleep stage variables and overall memory performance.

**Figure 7.**
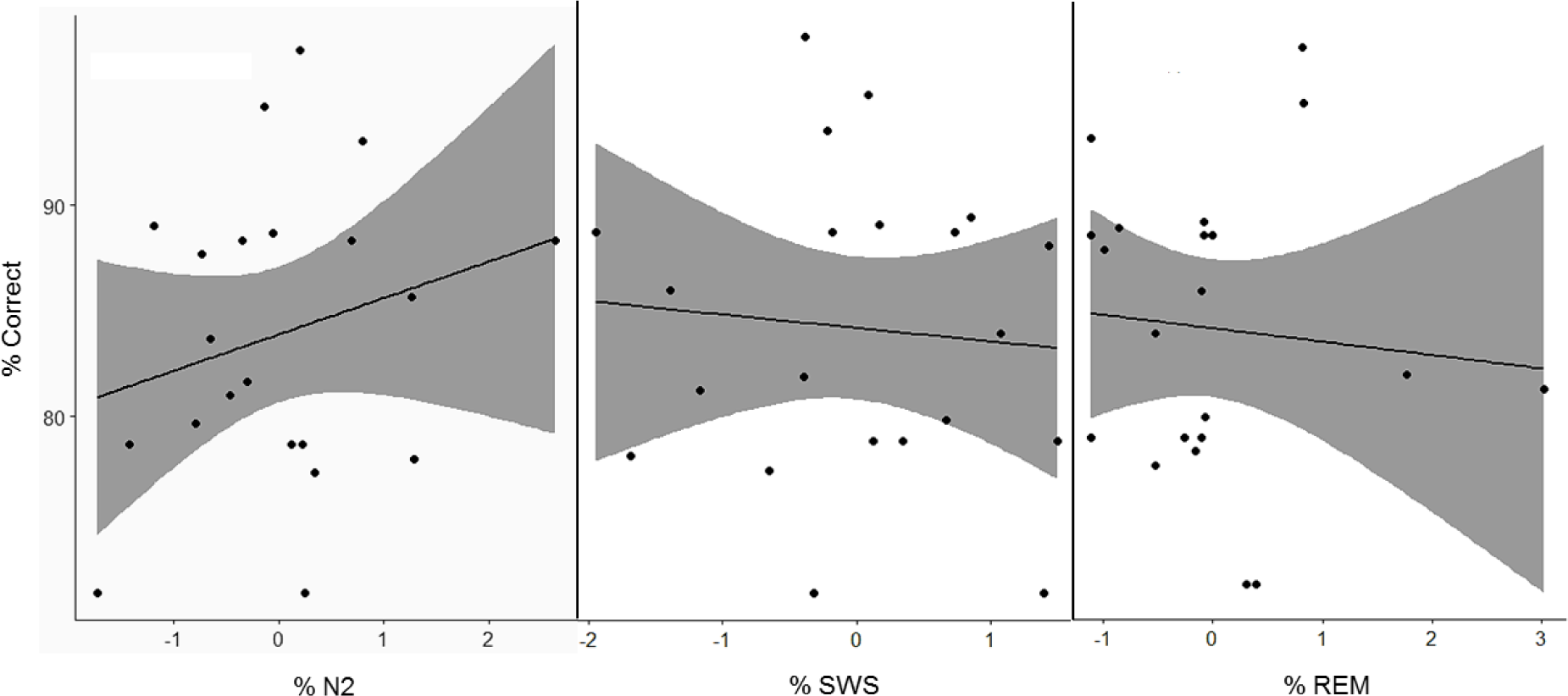
Scatterplots illustrating the relationship between the percentages of time spent in N2, SWS and REM on percentage of correct responses at delayed testing during the Sleep condition. Shaded space indicate the 95% confidence interval.

**Table 5.** Pearson correlations (n=21) between sleep stage variables and percentage of correct responses for negative, neutral and positive stimuli. The final row reports the relationship between the percentages of time spent in each sleep stage with overall memory performance.

|  | N2 | SWS | REM |
| --- | --- | --- | --- |
| Negative | 0.31 | -0.13 | -0.12 |
| Neutral | 0.08 | 0.02 | -0.006 |
| Positive | 0.27 | -0.12 | -0.11 |
| **Overall** | **0.24** | **-0.08** | **-0.09** |

## 5. Discussion

This study aimed to determine whether attentional state prior to encoding influences sleep-related memory consolidation for emotionally valenced, compared to neutral stimuli. Contrary to expectations, memory performance was significantly greater after a period of wake than after a 2hr sleep opportunity. Further, emotional valence did not appear to modulate memory, with no difference in memory performance between positive, negative and neutral stimuli. However, results indicated that memory was modulated by attentional state – as indexed by oscillatory alpha activity – and this effect was larger for the wake than the sleep condition. We interpret these findings in accordance with the inhibition-timing hypothesis (Klimesch et al., 2007), and proposals that memory consolidation benefits from the cyclic occurrence of SWS and REM (e.g. Giuditta, 2014).

### 5.1. Alpha, affective attention and emotional memory

Previous research utilising alpha as an index of attention and cortical inhibition have reported increases in alpha desynchronization to emotional stimuli (Uusberg et al., 2013). This is suggested to reflect a gating mechanism, preferentially processing biologically relevant information, while selectively filtering out less salient information (Aftanas et al., 2002; Uusberg et al., 2013). However, the present study found that enhanced alpha synchronisation prior to encoding resulted in greater memory performance at recall, which is consistent with models that posit alpha synchronisation plays an active role in global neural integration rather than passive neural idling (Basar, Shurmann, Basar-Eroglu & Karakas, 1997; Hanslmayr, Gross, Klimesch & Shapiro, 2011; Klimesch et al., 2007).

According to the inhibition-timing hypothesis (Klimesch et al., 2007), an increase in alpha synchronisation reflects the inhibition of bottom-up processing via attentional mechanisms. As such, the relationship between increased alpha synchronisation and greater memory consolidation might reflect a top-down attentional mechanism for sheltering information from external, task-irrelevant inputs (Hanslmayr et al., 2011). Further, there is strong evidence indicating that increased alpha power regulates the flow of information in the cortex via the synchronised firing of neuronal ensembles in occipital-parietal cortices (Klimesch et al., 2007; Schurmann & Basar, 2001). Cycles of neuronal excitability within these regions may act as a pulsed-inhibition gating mechanism, modulating the flow of task-relevant information to regions further downstream (i.e. entorhinal cortex and regions CA1/CA3) for long-term consolidation (Buzsaki, 1996). This interpretation is in line with evidence demonstrating that the phase of alpha oscillations modulates synaptic spiking, maximising stimulus detection and subsequent mnemonic processing (Canolty & Knight, 2010). For example, Khader and colleagues (2010) reported increased alpha synchronisation while participants engaged in internal mental processing tasks after encoding, which predicted enhanced performance on a delayed recall task. Thus, our findings build on the proposal that alpha oscillations serve as a gating mechanism, modulating memory encoding within a thalamo-neocortical-hippocampal network, opposed to the view that alpha synchronisation merely reflects cortical idling (e.g., Pfurtscheller et. al., 1996).

However, it is important to note the methodological differences between the current study and analogous research on the affective modulation of alpha. Aftanas et al. (2002) and Uusberg et al. (2013) measured event-related spectral perturbations (ERD) in alpha power, finding emotional stimuli to evoke greater alpha ERD than neutral images. While the ERD technique indexes changes in oscillatory activity on a millisecond timescale, it can disregard the importance that the overall brain state has on LTM formation (Hanslmayr, Staudigl & Fellner, 2012). Thus, naturally occurring fluctuations between alpha synchronisation and desynchronisation may promote enhanced cortical processing (Klimesch, 2012; Mathewson et al., 2011; Peterson & Voytek, 2017). This is consistent with our findings, where higher alpha power predicted the consolidation of images, supporting the idea that fluctuations in neuronal excitation promote memory formation (Fell & Axmacher, 2011). Future research indexing alpha ERD to individual stimuli should also measure rhythmic activity over extended periods of time, as done in this study, which could further elucidate how fluctuations between excitation and inhibition coordinate the formation of hippocampal-dependent memory traces (Hanslmayr et al., 2012).

### 5.2. Sleep and emotional memory consolidation

Based on research demonstrating a beneficial effect of sleep on memory consolidation generally (Rasch & Born, 2013; Schreiner & Rasch, 2014; Stickgold & Walker, 2013), and a sleep-related effect on emotional memory consolidation specifically (Groch et al., 2013; Lewis et al., 2011; Payne, Chambers & Kensinger, 2012; Payne et al., 2015; Wagner et al., 2001), we predicted that memory would be greater after a 2hr nap compared to wake, and that this effect would be accentuated for emotional compared to neutral stimuli. In contrast, memory degradation was greater after sleep compared to wake, coupled with no statistically significant difference in memory performance between negative, neutral, and positive stimuli.

Although offering many methodological advantages, such as controlling for circadian effects, an afternoon nap typically lacks REM sleep (Genzel, Spoormaker, Konrad & Dresler, 2015; Payne et al., 2015). Without REM to target neural representations of emotionally valenced items, memory for emotional stimuli may have weakened over time (Nishida, Pearsall, Buckner & Walker, 2009). This interpretation supports studies reporting a beneficial role of REM in promoting emotional plasticity, particularly through theta oscillations (Heib et al., 2015; Hutchison & Rathore, 2015). REM theta oscillations reflect coherent activity between the hippocampus and amygdala and are thought to preferentially strengthen emotionally tagged memory traces (Bennion, Payne & Kensinger, 2015; Hutchison & Rathore, 2015). As such, the effect of sleep reported here may be explained by a lack of REM theta activity, which preferentially strengthens memory traces of emotional stimuli after the synaptic downscaling induced by slow oscillations during SWS sleep (Hutchison & Rathore, 2015). This idea is in part supported by the small-to-moderate positive correlations between the percentage of time spent in N2 with memory for negative and positive but not neutral stimuli. N2 is dominated by theta activity and thalamic spindles and has been implicated in the consolidation of declarative and non-declarative (e.g., procedural) memory consolidation (for review: McDevitt, Krishnan, Bazhenov & Mednick, 2017). Similar to REM theta oscillations, theta activity during N2 may have supported the consolidation of emotional stimuli by facilitating the replay of encoded emotional memory traces between the prefrontal cortex and limbic regions (Bennion, Payne & Kensinger, 2015). By contrast, sleep spindles, which are argued to initiate synaptic plasticity via long-term potentiation (Cross, Kohler, Schlesewsky, Gaskell & Bornkessel-Schlesewsky, 2018; Staresina et al., 2015), may have facilitated the transfer of hippocampally dependent (emotional) memory traces with neocortical LTM networks, which is consistent with research reporting that pharmacologically increasing sleep spindles enhances emotional memory (Kaestner, Wixted & Mednick, 2013).

### 5.3. Alpha oscillations, sleep and memory consolidation

This study provides preliminary evidence that attention and sleep modulate memory consolidation in distinct ways. Greater attention prior to encoding appears to be important for preserving memory during wakefulness, and that a 2hr sleep opportunity – without sufficient REM – may transiently shelter memory from the external inputs of wakefulness. One possible explanation for the distinction between wake and sleep findings comes from consideration of the interplay of alpha activity with that of theta. Theta rhythms during wake have been associated with enhanced memory formation, wherein after learning emotional stimuli, theta activity has been reported to synchronise between the hippocampus and temporal cortex, reflecting hippocampal-cortical communication (Brenner et al., 2014; Hutchison & Rathore, 2015; Vertes, 2005). During sleep, theta is predominantly evident during REM – a stage limited in the present study by the use of a nap paradigm – which is characterised by an increased production of acetylcholine, an important factor for neuroplasticity (Heib et al., 2015; (Scheffzuk et al., 2011). During the sleep and wake conditions, synchronous alpha activity may have reflected top-down control critical for preventing interference of recently established memory traces, reducing irrelevant information from reaching enhanced processing (Khader et al., 2010; Klimesch et al., 2007). Following encoding, during the wake condition, theta activity may have strengthened memory traces between the hippocampus and amygdala (Heib et al., 2015). However, as the sleep period in this study contained minimal REM, memory traces would be weakened due to the lack of theta activity, leading to poorer memory performance at delayed recall (Hutchison & Rathore, 2015).

The finding that greater pre-encoding alpha synchronisation protected memory consolidation after wake compared to sleep can be interpreted within the context of the synaptic homeostasis hypothesis (SHY; Tononi & Cirelli, 2014). According to SHY, sleep facilitates the downscaling of synaptic weight accumulated during wake to a baseline level that is homeostatically sustainable, a process performed by slow oscillations during SWS (Mascetti et al., 2013). This process of synaptic renormalisation desaturates the capacity to encode new information during subsequent wake periods by decreasing neuronal excitability, which improves the signal-to-noise ratio in the reactivation of encoded memory traces (Tononi & Cirelli, 2012). It is argued that one function of alpha synchronisation is to prevent competing memory traces from being concurrently activated during memory retrieval (Hanslmayr et al., 2012). From this perspective, the wake condition – having not had the benefits of sleep-based synaptic downscaling – may have required greater internally oriented processing to distinguish between new and old items. In contrast, the selective downscaling of synaptic weights during sleep in the nap condition may have fine-tuned connections between occipital-parietal cortices and neuronal assemblies further downstream, facilitating quick and efficient memory retrieval. Future research should directly test whether the occurrence of SOs during SWS interact with alpha activity during encoding on subsequent memory performance.

Taken together, theta oscillations – present during N2 and markedly increased during REM – may serve complementary roles in the emotional modulation of LTM, and alpha, as a physiological marker of attention and cortical inhibition, may facilitate this process at encoding. However, future research is required to determine the interactive effect that alpha and theta activity have on the emotional modulation of LTM across periods of sleep versus wake. Such research would shed further light on the factors underpinning the attentional modulation of sleep-related memory consolidation, which may lead to a greater understanding of disorders in which attentional deficits to emotional information modulate sleep-related learning (Baran, Pace-Schott, Ericson & Spencer, 2012). For example, Prehn-Kristensen et al. (2013) found that sleep-facilitated consolidation of emotionally valenced stimuli was impaired in children with attention-deficit-hyperactivity disorder, but not healthy controls, highlighting the role attention may play in sleep and the emotional modulation of LTM.

### 5.4. Limitations and strengths of current study

As discussed, one major limitation of the current study was minimal REM sleep occurring during daytime naps, as well as large variability in the time participants slept. These effects make it difficult to establish a true effect of sleep. Future research using a nocturnal half-night paradigm would involve presenting participants emotional stimuli before either a SWS or REM-rich sleep interval (Groch et al., 2013). Nocturnal half-night paradigms account for natural human nocturnal sleep architecture, and would allow for the systematic investigation between alpha activity at encoding and memory consolidation during SWS and REM.

The adoption of a nocturnal half-night paradigm would be complemented by the use of individual valence reports instead of normalised ratings. While the majority of research utilising the IAPS has relied on normalised ratings (Bradley & Lang, 1994; Lang & Bradley, 2007), there may be large inter-individual variability in the way subjects perceive stimuli along the dimensions of valence and arousal (Backs, Silva & Han, 2005). Quantifying emotional memory based on individual self-report ratings may increase model sensitivity and the probability of detecting emotion-enhanced memory effects, similar to previous emotional memory research (for review: Talmi, 2013).

Finally, while averaging alpha power over the course of the CCPT-V enabled quantification of a cortical index of sustained attention, it did not allow for direct observation of event-related changes in the power spectra. While this method yields distinct strengths, it limits the ability to establish links between changes in alpha power at the point of stimulus presentation, making it difficult to infer the relationship between alpha (de)synchronisation and the consolidation of stimulus-specific characteristics. Despite these limitations, the adjustment of the alpha range according to participants’ IAF yields greater reliability over utilising the standard 8–12 Hz range. The distribution of alpha-band activity shows substantial variation across individuals, and has been associated with inter-individual differences in information processing and general intelligence (Klimesch, 1997; 1999). Further, anchoring frequency band windows in relation to the IAF as opposed to classically defined bandwidths has been shown to increase sensitivity to band-specific oscillatory dynamics (Doppelmayr, Klimesch, Pachinger, & Ripper, 1998). From this perspective, the effects of alpha reported here are unlikely to be due to overlap with other frequency band activity (e.g. beta, 13–30 Hz; for a discussion on alpha/beta activity and long-term memory, see Hanslmayr et al., 2012).

## 6. Conclusions

This study demonstrated that attention, as indexed by alpha oscillatory activity, modulates memory consolidation during wake relative to an equivalent period of sleep. These results address a neglected area of research into the sleep-dependent memory consolidation hypothesis: effects of sleep versus wake on the consolidation of various types of memory are often assessed in isolation to memory encoding and attentional state, limiting the ability to separate such factors from sleep-facilitated consolidation on subsequent memory. From this perspective, alpha oscillatory activity provides a valuable pathway for investigating the neural correlates of attention and sleep-facilitated memory consolidation. However, in line with previous accounts, we posit that REM oscillatory theta activity may underlie post-encoding processes involved in memory consolidation, particularly for emotionally valenced information. Future research should directly measure the combined effect of alpha at encoding, and theta during consolidation, across periods of wake and sleep. This will provide a valuable basis for future research investigating brain mechanisms involved in the interactive effect of attention and sleep on long-term memory consolidation.

## Acknowledgements

We thank Professor Ina Bornkessel-Schlesewsky for helpful comments on an earlier version of this manuscript, and Alex Chatburn for valuable discussion. The authors declare no competing financial interests.

